# MERFISHEYES: Web-based Visualization and Sharing Platform for Single-cell and Single-molecule Spatial Transcriptomics

**DOI:** 10.64898/2026.09.20.753029

**Authors:** Ignatius Jenie, Evan Mishkin, Elvis Smith, Mariah Kenney, Ivan Cao-Berg, Roy Maimon, Alexander J. Ropelewski, Don W. Cleveland, Bogdan Bintu

## Abstract

Interactive exploration and sharing of hybridization-based spatial transcriptomics datasets remain limited by large file sizes, specialized software, and computationally intensive preprocessing, restricting rapid access to publicly available resources such as the Brain Image Library. Here we present MERFISHEYES (https://merfisheyes.com), a browser-based platform for interactive visualization of single-cell and single-molecule spatial transcriptomics data across commercial and emerging high-throughput platforms. MERFISHEYES enables marker gene exploration, cross-dataset and cross-species comparison, visualization of subcellular RNA localization, and collaborative sharing through browser-accessible links generated directly from drag-and-drop datasets. We use MERFISHEYES to enable browser-based visualization of all 158 spatial transcriptomics datasets in the Brain Image Library and demonstrate its capabilities in handling high-throughput 3D data with genome-scale imputation of gene expression and DNA accessibility. MERFISHEYES provides a scalable framework for rapid exploration, sharing, and collaborative analysis of spatial transcriptomics data.

**Highlights:**

- MERFISHEYES is an online web-based visualizer for spatial transcriptomics data that requires no user preprocessing, installation, or hosting of data
- All 158 datasets from the Brain Image Library are integrated into MERFISHEYES allowing for cross dataset comparisons
- Supports single-molecule visualization of more than 150 million transcripts in 3D
- Supports link-based data sharing for easy access and collaboration of spatial transcriptomics data

## Introduction

Spatial transcriptomics encompasses a collection of growing technologies which enable characterization of cellular diversity and spatial distribution of gene expression within intact tissue architecture^1–5^. A subclass of spatial transcriptomics are based on repeated cycles of hybridization and imaging, including methods such as Multiplexed Error-Robust Fluorescence In Situ Hybridization (MERFISH)^6,7^, Seq-FISH^8^, and others^9–14^. These methods complement previously established sequencing-based methods such as Slide-seq^15,16^, Stereo-Seq^17^ and others^18–22^. Hybridization approaches allow for both the highest detection efficiency of targeted genes and the highest spatial resolution by localizing mRNA molecules within 3D subcellular environment^23^.

Over the past five years, commercial platforms such as Vizgen’s MERSCOPE^24^ and 10X’s Xenium^25^, have facilitated a broad range of application of these technologies^4^. The Brain Image Library, a repository for large brain image datasets, has amassed 158 spatial transcriptomics datasets totaling to 132 million profiled cells. However, the resulting datasets, each reporting levels of RNAs encoded by ∼600 different genes out of the more than 20,000 protein coding genes in the human genome, are difficult to explore and compare. A single experiment typically localizes hundreds to thousands of molecules within singular cells and contains hundreds of thousands of individual cells^26–28^. Recent advances in high throughput imaging, which enables 3D reconstruction of entire human organs^29^, whole-transcriptome coverage^30^, and high-resolution continuous 3D imaging^29,31,32^ further compound the size and complexity of these datasets.

Visualizers developed by commercial vendors^33–37^ allow for data exploration but are closed source and only accept data generated on their specific platforms, inhibiting comparisons across the different datasets available. Conversely, open-source viewers have the flexibility of loading all existing datasets but require extensive computational expertise to deploy and host server infrastructure^38^. Both commercial and open-source visualizers share a critical limitation: they are built around transcripts preassigned to segmented cells and offer little or no support for visualizing individual transcripts, so the subcellular signal is either aggregated or accessible only via subsampling^33,34,38^. No existing tool allows opening both single-molecule and single-cell spatial transcriptomics data in web-based interface without installing software or running infrastructure; none preserve transcript-level resolution for more than 150 million molecules, and none allow datasets to be compared against one another within a common cell type taxonomy.

We had previously built custom versions of a browser-based viewer for three studies: a comprehensive survey of olfactory receptor expression^39^, a 3D multiome-imputed spatiotemporal atlas of the zebrafish embryo^31^, and a 3D atlas of the developing human heart^29^. Although these initial implementations were designed and optimized specifically for serving the corresponding datasets, they helped establish the foundation for a generalized platform.

Here we present MERFISHEYES (https://merfisheyes.com), a browser-based platform to explore, compare, and share single-cell and single-molecule spatial transcriptomics data. Datasets are uploaded via drag-and-drop, processed locally in the browser and uploaded to return a shareable browser link. This pipeline requires no local installation, manual preprocessing, server management, or specialized computational expertise. Data can be examined at two distinct resolutions: 1) the single-cell level, which enables exploration of the spatial organization of different cell-types, and 2) the single-molecule level, which retains the subcellular signal within nuclei, soma and cellular processes that single-cell-level analysis post-segmentation discards. At the single-molecule level, MERFISHEYES renders all molecules simultaneously without subsampling, making full-resolution, transcript-level data explorable from the browser for the first time. For unpublished data, these capabilities facilitate quality inspection and data sharing among collaborators via explorable web-links, as previously achieved for genetic and epigenetic data^40-42^.

MERFISHEYES also serves as a community resource. It is coupled to the Brain Image Library (BIL)^43^, enabling the visualization of all previously deposited spatial transcriptomics data. We have mapped these datasets to a common cell-type taxonomy so that matching populations can be cross-compared. Since the corpus is already accessible via MERFISHEYES and cell type definition has been standardized, any existing datasets can be easily compared against each other, spanning various imaging platforms, treatments, and species. Overall, MERFISHEYES provides both an explorable platform and a comprehensive resource for neuroscience.

## Results

### Drag and drop in-browser visualization of single-cell and single-molecule spatial transcriptomics data

Spatial transcriptomics data is inherently organized at two levels of resolution: single-cell and single-molecule. First, at the single-cell level, the number of RNA transcripts measured in each segmented cell is quantified across genes^4^. The table of transcript counts across genes and cells, similar to single-cell (or single nucleus) RNA sequencing data, allows for transcriptionally defining cell-types and representing single cells in both transcriptional space using UMAP embeddings^44^ and spatially within the larger tissues/organ architecture. Second, at the single-molecule level, spatial transcriptomics localizes individual RNA molecules subcellularly within individual cells allowing for categorizing genes associated within different cellular compartments (nucleus, cytoplasm, processes, etc.).

We built MERFISHEYES for this dual-mode exploration of both single-cell and single-molecule data types, without requiring installation, preprocessing, or computational expertise (**Fig. 1a-c**). The visualizer is built around a browser-native architecture (Next.js) in which data parsing and rendering are performed entirely on the user’s device. Outputs from the most used commercial platforms (MERSCOPE, Xenium) and the more general AnnData files (.h5ad) for single-cell data, and tabular formats (.parquet, .csv) for single-molecule data, are parsed via open-source libraries (h5wasm, anndata.js, hyparquet, papaparse) into a common internal representation (Methods; **Fig. 1d**). Parsing happens in seconds: a 382,000-cell × 275-gene .h5ad loads in 30 seconds, a 258,000-cell × 541-gene Xenium run in 7.5 seconds, an 89,000-cell × 649-gene MERSCOPE run in 18 seconds, and single-molecule datasets of up to 32 million transcripts can render in under a minute (**Supplementary Fig**. **1a-d**). Current limitation of data size is bounded by the browser which require uploaded files to be under 2 GB.

**Figure 1.**
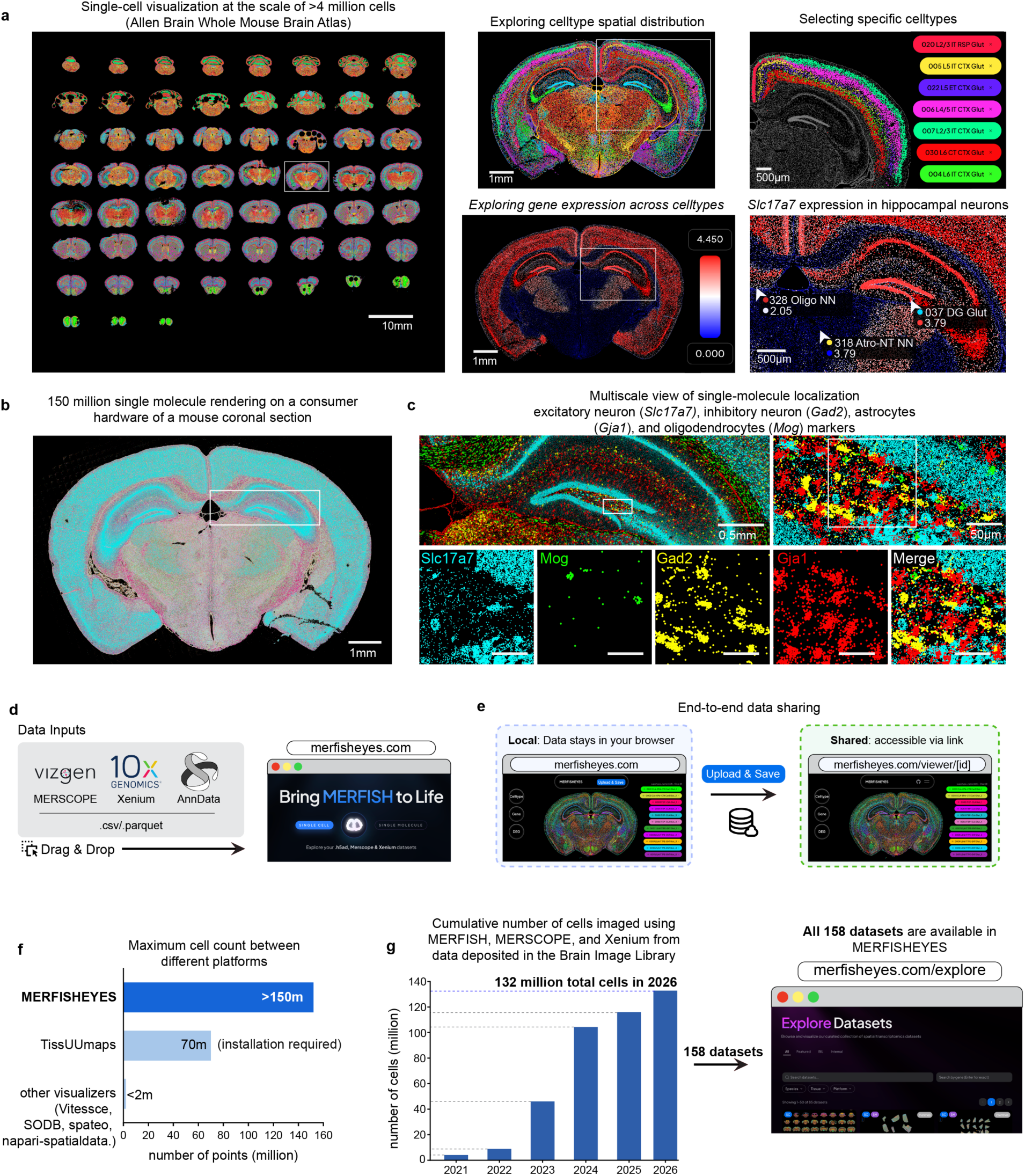
Browser-native visualization of single-cell and single-molecule spatial transcriptomics. **a**, Single-cell visualization of the Allen Institute Whole Mouse Brain Atlas^47^ (∼4 million cells) in MERFISHEYES. Left, all coronal sections rendered at once, cells colored by subclass-level cell type; scale bar, 10 mm. Top middle, the section boxed at left, cells colored by subclass; scale bar, 1 mm. Top right, magnification of the boxed neocortical region, with seven glutamatergic subclasses selected (020 L2/3 IT RSP Glut, 005 L5 IT CTX Glut, 022 L5 ET CTX Glut, 006 L4/5 IT CTX Glut, 007 L2/3 IT CTX Glut, 030 L6 CT CTX Glut, 004 L6 IT CTX Glut) and all other cells in grey; scale bar, 500 µm. Bottom middle, the same section with cells colored by *Slc17a7* expression (log-normalized; blue, 0; red, 4.45); scale bar, 1 mm. Bottom right, magnification of the boxed hippocampal region showing *Slc17a7* enrichment in the dentate gyrus, CA1 and CA3. Hovering over a cell opens a tooltip with its cell type and expression value (three examples shown: 328 Oligo NN, 037 DG Glut, 318 Astro-NT NN); scale bar, 500 µm. **b**, Single-molecule visualization of the same coronal section, showing all ∼150 million detected transcripts at once, without subsampling or aggregation; scale bar, 1 mm. **c**, Single-molecule distribution of four cell-type markers within the region boxed in **b**: *Slc17a7* (excitatory neurons, cyan), *Mog* (oligodendrocytes, green), *Gad2* (inhibitory neurons, yellow) and *Gja1* (astrocytes, red). Top left, merged view of the hippocampal region; scale bar, 0.5 mm. Top right, magnification of the boxed area; scale bar, 50 µm. Bottom, the region boxed in the top-right panel, shown for each marker separately and merged; scale bars, 30 µm. **d**, Supported inputs. Vizgen MERSCOPE and 10x Genomics Xenium output folders, AnnData (.h5ad) files and single-molecule tables (.csv, .parquet) are loaded by drag-and-drop at merfisheyes.com and parsed on the user’s device; nothing is uploaded or preprocessed before visualization. **e**, Data sharing. Left, a dataset opened locally stays in the browser. Right, on Upload & Save the dataset is compressed client-side into a chunked streamable format, transferred to cloud object storage and returned as a persistent link (merfisheyes.com/viewer/[id]) that reopens the same view on any device. **f**, Maximum number of points rendered in a single view by browser-based spatial transcriptomics visualizers: MERFISHEYES, >150 million; TissUUmaps, 70 million; all other tools tested, <2 million. Values are the largest point count each tool displayed without subsampling on a MacBook Air M3; tools, versions and the benchmark protocol are listed in Table 1 and Methods. **g**, MERFISHEYES as the Brain Image Library viewer. Left, cumulative number of cells imaged by MERFISH, MERSCOPE and Xenium in the 158 datasets deposited in the Brain Image Library, 2021-2026 (132 million cells in total). Right, all 158 datasets can be browsed and filtered by species, platform and gene at merfisheyes.com/explore.

**Table 1.**
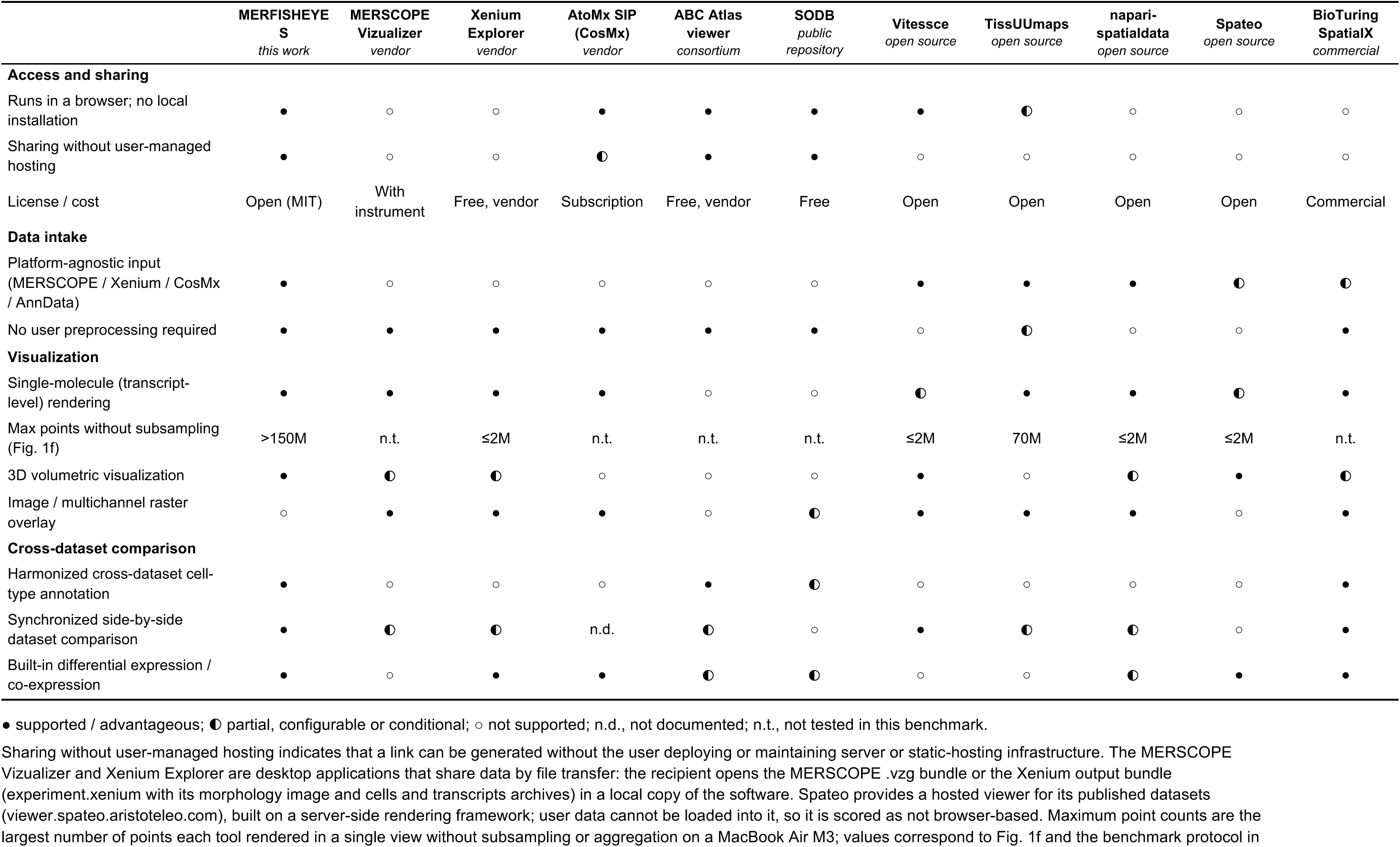
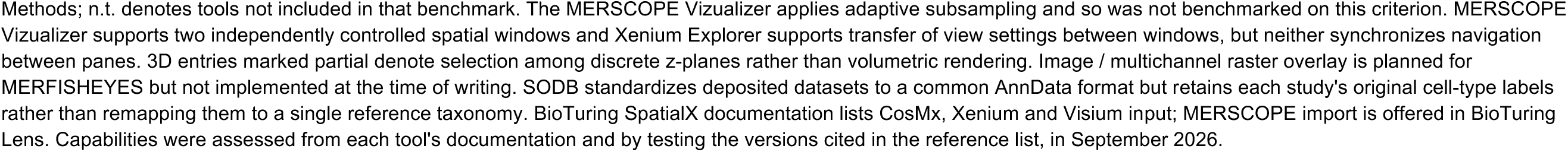
Capability comparison of MERFISHEYES with existing spatial transcriptomics visualization tools.

For datasets beyond these limits, MERFISHEYES automatically performs the conversion server-side through the same drag-and-drop flow. In our benchmarks this allows an order of magnitude increase in data size upload: a 3-million-cell MERFISH .h5ad (11 GB decompressed) is converted and visualizable in about an hour, a whole-transcriptome Xenium run (717,576 cells × 27,104 genes) in 44 minutes, and a 669-million-transcript single-molecule dataset in 13 minutes. Jobs that exhaust a memory tier are retried automatically on larger machines (16 as default, 32, then 64 GB), and a typical dataset costs between $0.01-0.30 USD of compute (Methods; **Supplementary Fig. 1a-d**).

The same architecture supports web-based sharing. Once a dataset is loaded, users can upload it directly from the browser and receive a shareable link to a cloud-hosted viewer (**Fig. 1e**). Before any data leaves the device, the dataset is compressed into a lightweight streamable format, so only a compact representation is transferred to cloud storage (Methods). The resulting link works for unpublished and published data alike: collaborators or reviewers can explore an unpublished dataset interactively without downloading files or installing software, and a published dataset can be displayed openly as a public atlas.

### GPU-accelerated rendering supports visualizing more than 150 million single-cells or single-transcripts

Visualizing spatial transcriptomics data requires rendering up to millions of cells and tens of millions of single-molecules: a task that exceeds the capacity of most existing viewers^33–35,37,38^, which typically subsample or aggregate the data before display. MERFISHEYES addresses this through a rendering layer built on three.js and WebGL, rendering more than 150 million points in a single view on a standard laptop (**Fig. 1b,f).** This rendering capacity more than doubles that of the next best browser-based visualizer (TissUUmaps^45^, 70 million points) and greatly exceeds that of widely used tools^32–34,38,46^ (**Fig. 1f**). Combined with the browser-native parsing described above, MERFISHEYES displays full single-molecule spatial transcriptomics datasets at native resolution without subsampling.

### Interactive single-cell exploration of multi-million-cell spatial atlases and single-molecule visualization at subcellular resolution

MERFISHEYES enables single-cell visualization of the largest spatial transcriptomics data directly in the browser. We demonstrated this capability by loading in the entire Mouse Brain Atlas compiled by the Allen Institute^47^ consisting of ∼4 million cells (**Fig. 1a**). Due to the comprehensive nature of this brain-wide murine dataset we made it easily accessible at *merfisheyes.com/explore* for easy comparison with other datasets. In MERFISHEYES cells are colored by transcriptionally defined cell type identity at 4 different resolutions^47^ (class, subclass, supertype and cluster level). Taking subclass level, as an example, the platform allows for double-click selecting and deselecting specific cell types. While each cell type is defined based on the single-cell transcriptional profiles, MERFISHEYES reveals which cell type are constrained to well defined anatomical structures, such as the distinct layered architecture of the neocortex (**Fig .1a**, top right) and which cell types are more broadly distributed throughout the brain, such as microglia (**Supplementary Fig. 1e**). For each targeted gene, single-cell expression can also be explored. For example, the excitatory neuron marker *Slc17a7* shows strong enrichment across hippocampal subfields of DG, CA1, and CA3 neurons (**Fig. 1a**, bottom right). Zooming and panning support inspection across anatomical scales, hovering displays per-cell metadata, and double-clicking isolates a selected cell type population.

On the same coronal section where we displayed cell type distribution using single-cell mode, we demonstrate the single-molecule mode by displaying all 150 million transcripts detected (**Fig. 1b**). This eliminates the data reduction or aggregation typically required by other visualization tools, preserving the full spatial information at single-molecule resolution. We interactively sub select four canonical cell type markers, *Mog* (oligodendrocytes), *Gad2* (inhibitory neurons), *Slc17a7* (excitatory neurons), and *Gja1* (astrocytes), capturing the morphology of major cell populations (**Fig. 1c**). This visualization enables inspection of transcript positions down to the subcellular scale, resolving where individual molecules sit relative to cell boundaries.

### MERFISHEYES visualizes the entire Brain Image Library spatial transcriptomics data

The Brain Image Library (BIL)^43^ is a repository for large brain image datasets that has amassed 158 spatial transcriptomics datasets since 2021, totaling to 132 million cells imaged (**Fig. 1g**). The single-molecule measurements associated with these data and most of the single-cell datasets are not available for exploration. Accessing these datasets requires downloading files locally or running analyses on BIL’s compute infrastructure. Furthermore, datasets carry inconsistent cell-type annotations or lack searchable metadata of gene panels profiled, impeding cross-dataset comparisons.

We made all 158 datasets explorable in MERFISHEYES under the *Explore* tab (https://merfisheyes.com/explore). This makes MERFISHEYES the first platform to provide interactive single-molecule visualization of over a hundred brain datasets spanning multiple species, including recent large scale Brain Initiative effort covering the entire mouse brain and basal ganglia atlases of human, macaque and marmoset. Datasets can be explored and filtered by species, platform, and gene. The gene filter is particularly useful because spatial panels typically cover only a few hundred to a thousand genes, so researchers can confirm a dataset contains their genes of interest before exploring it, rather than downloading each dataset and realizing the gene of interest is not present. This reduces the time to test new hypotheses against published data.

### Cell type standardization for all BIL data using MapMyCells

The datasets deposited in BIL, which span different imaging platforms, gene panels and experimental treatments, do not have a coherent cell type definition or lack cell type definition altogether. To facilitate comparison between different datasets, we mapped all mouse, non-human primate, and human datasets to a single reference taxonomy using MapMyCells (MMC)^48^, a tool that labels each cell by correlating its expression profile to Allen Institute’s reference single-cell taxonomy^47,49^. This taxonomy is hierarchical spanning class, subclass, supertype and cluster level annotations in increasing order of resolution.

Our mapping reveals that automated cell type mapping works best for adult and post-natal mouse samples, and reveal less accurate mapping in neonatal mouse samples as well as human and non-human primate samples at high cluster-level resolution (**Supplementary Fig. 2a**). These results suggest that improved single-cell reference atlases are needed for neonatal mouse and human/non-human primate datasets. We provide the overall mean per-cluster correlation for each dataset when searching the datasets on merfisheyes.com/explore as measure of the cell annotation quality.

With these standardized cell type annotations, MERFISHEYES enables a split-synchronized view of two datasets allowing direct comparison (**Fig. 2a**). Taking a 4-week murine brain data^50^ from BIL as an example, and synchronizing it with the 8-week murine brain ABC atlas as a reference, we observed a consistent spatial distribution of matching cell types between the transferred annotation and the reference. When focusing on the *caudoputamen*, we recapitulated both D1 and D2 subclass-level medium spiny neurons (MSNs) restricted to the striatum (**Fig. 2b**). At the finest, cluster-level resolution, we observed the dorsolateral-ventromedial regional organization appearing consistently in both datasets across both direct and indirect pathway MSNs (0951 and 0982 enriched in the dorsolateral and 0950 and 0981 enriched in the ventromedial regions) (**Fig. 2c**). This regional segregation is consistent with the topographic sensorimotor and associative *corticostriatal* projections to the striatum^51,52^. Together, this exploration confirms the quality of the dataset and cell type annotation through direct one-to-one comparison.

**Figure 2.**
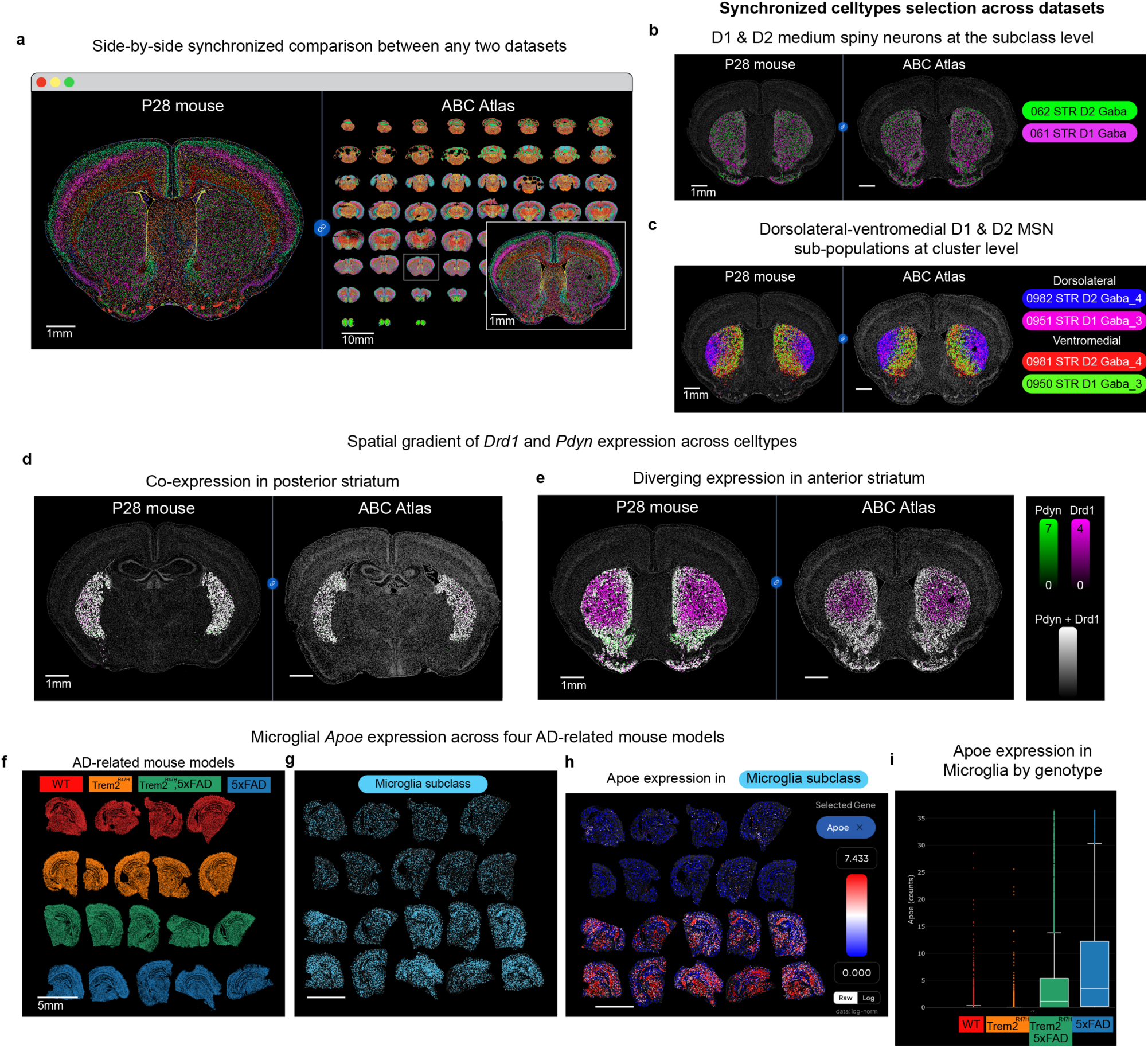
MERFISHEYES enables direct comparison across Brain Image Library datasets. **a**, Split-synchronized view of two independently generated datasets after both were mapped to the Allen Institute Whole Mouse Brain single cell reference taxonomy with MapMyCells. Left, a P28 developing mouse brain section^50^;right, the ABC atlas^36^. Cells are colored by subclass; cell-type and gene selections are synchronized between the two panes. Scale bars, 1 mm (P28 section and ABC inset) and 10 mm (ABC overview). **b**, D1 and D2 medium spiny neuron subclasses (061 STR D1 Gaba, magenta; 062 STR D2 Gaba, green) in the caudoputamen of the P28 dataset (left) and the ABC atlas (right); all other cells, grey. Scale bars, 1 mm. **c**, The same comparison at cluster level, separating dorsolateral (0982 STR D2 Gaba_4, blue; 0951 STR D1 Gaba_3, magenta) and ventromedial (0981 STR D2 Gaba_4, red; 0950 STR D1 Gaba_3, green) sub-populations in both datasets. Scale bars, 1 mm. **d**,**e**, Co-expression of *Pdyn* (green) and *Drd1* (magenta) within D1 clusters in anterior (**d**) and posterior (**e**) striatal sections, shown for the P28 dataset (left; anterior section^53^, posterior section^50^) and the ABC atlas (right). Colour scales are raw counts (*Pdyn*, 0-7; *Drd1*, 0-4); cells expressing both genes appear white. Scale bars, 1 mm. **f**, Coronal sections from four 12-month-old genotypes (Johnston et al., 2025; Brain Image Library ace-ear-nap^55^): wild type (red, 4 sections), Trem2R47H (orange, 4 sections), Trem2R47H;5xFAD (green, 5 sections) and 5xFAD (blue, 5 sections). All cells shown, colored by genotype. Scale bar, 5 mm. **g**, The same sections with only the microglia subclass displayed (cyan); all other cells, grey. Scale bar as in **f**. **h**, Microglia colored by *Apoe* expression (raw counts; blue, 0; red, 7.433). Scale bar as in **f**. **i**, *Apoe* counts in microglia by genotype.

### Identifying differentially expressed genes within cell types

MERFISHEYES enables identifying differentially expressed genes (DEG) and co-expression analysis across cell types. The in-browser analysis identified *Drd1* and *Pdyn* as DEGs distinguishing D1 from D2 MSN subclasses (**Supplementary Fig. 2b**) and their co-expression within D1 clusters was represented. While both genes co-localized frequently in the D1 neurons of the posterior striatum^53^ (**Fig. 2d**), anterior sections showed a diverging expression patterns between *Pdyn* and *Drd1* (**Fig. 2e**). This regional variation persisted even at the cluster level-the finest resolution available in the reference taxonomy-and is recapitulated across both datasets (**Supplementary Fig. 2c**). This exploration reveals heterogeneity in *Pdyn* expression within D1 MSNs that current annotations do not capture *Pdyn* encodes *dynorphin*, which inhibits striatal dopamine release and shapes reward-related behavior^54^, implying that graded expression across the D1 population may be functionally consequential.

### Application for exploring gene differences across mouse models

Beyond comparing cell types, MERFISHEYES supports visual and quantitative analysis of gene expression within a given cell type across experimental conditions, supporting disease-association analyses. We demonstrate this capability using a BIL dataset^55^ containing a cohort of four mouse genotypes (12 month-old): wild-type (WT), 5xFAD (transgenic amyloid model), Trem2^R47H^ (an Alzheimer’s Disease risk-variant knock-in), and Trem2^R47H^;5xFAD (**Fig. 2f**). We examined *Apoe*, a disease-associated microglia marker that is upregulated with amyloid pathology in the 5xFAD model^56^. Restricting the analysis to the microglia subclass, MERFISHEYES showed higher *Apoe* expression in the 5xFAD and Trem2^R47H^;5xFAD samples than in the WT and Trem2^R47H^ controls (**Fig. 2g-i**). The same increase was evident spatially (**Fig. 2h**): certain regions, such as the corpus callosum, contained a higher proportion of Apoe-high microglia. These analyses show that MERFISHEYES can locate and quantify cell-type-specific expression changes across disease models.

### Subcellular localization of single mRNA molecules

Beyond cell-level analyses, MERFISHEYES preserves single-molecule resolution, retaining the subcellular localization of transcripts in nucleus, cell body, and cellular processes that standard cell segmentation pipelines typically discard^57,58^. Standard cell segmentation, which relies on nuclear or membrane staining, labels each RNA transcript as “assigned” if within the computed segmentation mask of each cell or as “unassigned” if it lies outside segmentation boundaries. The single-molecule view enables visualization of both “assigned” and “unassigned” transcripts, revealing additional layers of information that would otherwise be discarded, including cell morphology differences across cell-types, transcript localization within cells and segmentation artefacts including transcript misassignment to neighboring cells and high levels of unassigned transcripts.

Collectively, the RNA transcripts within each cell approximate the extent of the cell soma and nearby processes allowing for identifying morphological differences between cell-types. A recent study from our group reported that striatal cholinergic (ChAT⁺) interneurons are among the largest-bodied striatal neurons, substantially larger than D1 medium spiny neurons in human^59^. We extended this comparison to rhesus macaque^60^ and mouse^50^ by clustering ChAT and Drd1 transcripts, which mark ChAT⁺ interneurons and D1 medium spiny neurons respectively (**Fig. 3a**). In both species, ChAT⁺ somas spanned ∼2× the diameter of D1 MSN somas (**Fig. 3b**), indicating that the cholinergic-to-D1 soma-size ratio is conserved across species.

**Figure 3.**
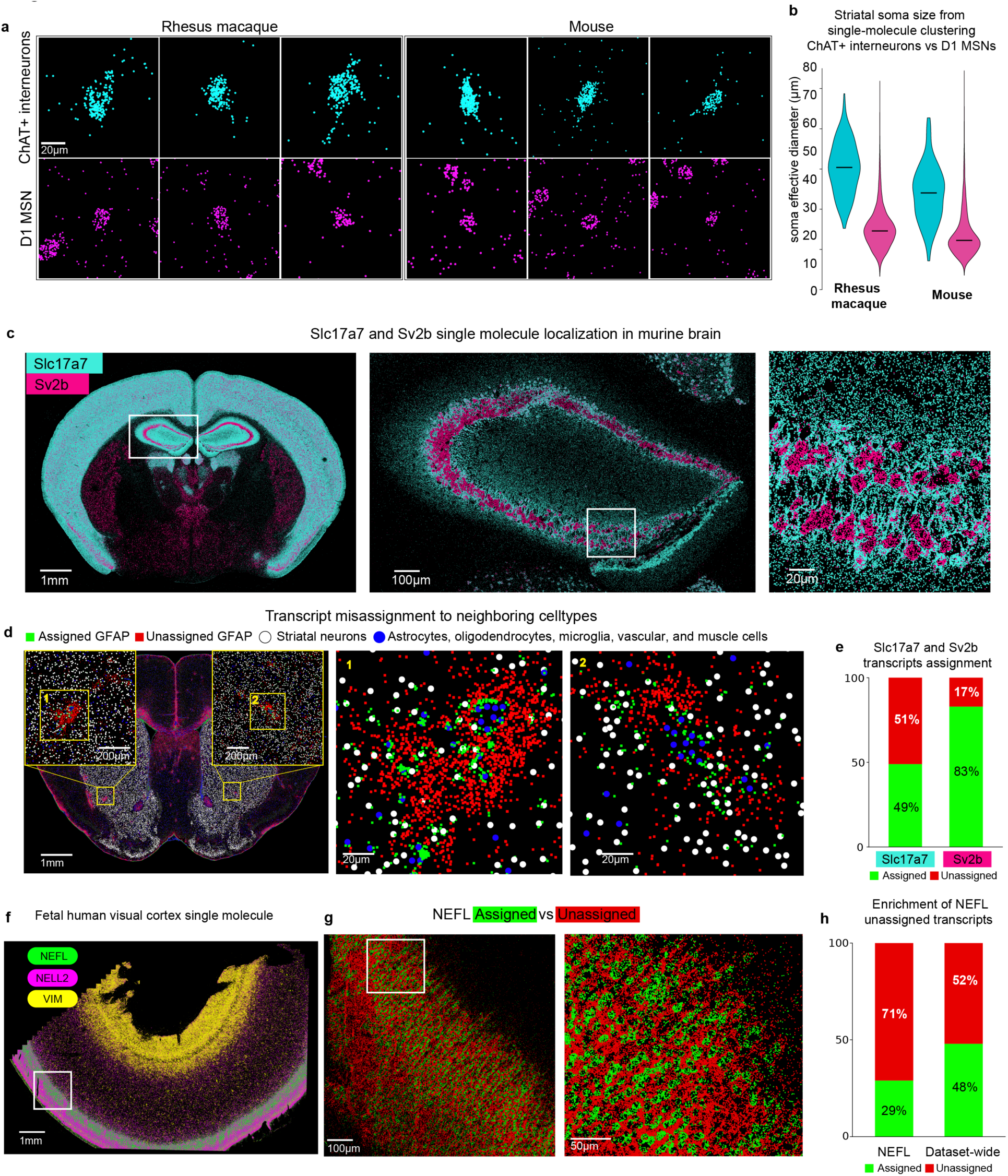
Single-molecule visualization resolves transcripts that cell segmentation discards. **a**, Soma reconstructed from transcript positions alone. *ChAT* transcripts (cyan, top row) mark cholinergic interneurons and *Drd1* transcripts (magenta, bottom row) mark D1 medium spiny neurons in striatal sections from rhesus macaque^60^ (left, three examples of each) and mouse^53^ (right, three examples of each). Scale bars, 10 µm. **b**, Effective soma diameter of *ChAT*+ interneurons (cyan) and D1 medium spiny neurons (magenta) in each species, from transcript clusters identified with DBSCAN. Center line, median; n = 178 ChAT+ and 10,152 D1 somata in macaque, and 202 ChAT+ and 6,931 D1 somata in mouse. One-sided Mann-Whitney U test (ChAT+ > D1): *P* = 2.0 × 10^−101^ (macaque) and 1.9 × 10^−78^ (mouse); ratio of medians, 2.08 (macaque) and 1.96 (mouse). **c**, *Slc17a7* (cyan) and *Sv2b* (magenta) transcripts in a mouse brain coronal section^75^. Left, whole section; scale bar, 1 mm. Middle, magnification of the boxed hippocampal region, with *Sv2b* confined to the pyramidal cell layer and *Slc17a7* extending into the surrounding neuropil; scale bar, 100 µm. Right, magnification of the boxed area; scale bar, 20 µm. **d**, Misassignment of *Gfap* transcripts to neighbouring neurons in the mouse striatum^53^. Assigned *Gfap* transcripts, green; unassigned, red; centroids of striatal neurons, white circles; centroids of astrocytes, oligodendrocytes, microglia, vascular and muscle cells, blue circles. Left, whole section with two boxed regions shown as insets; scale bar, 500 µm. Middle and right, regions 1 and 2 at higher magnification; scale bars, 20 µm. **e**, Fraction of *Slc17a7* and *Sv2b* transcripts assigned to a segmented cell (green) or unassigned (red) across the whole section. n = 1.8 million *Slc17a7* and 147.8 thousand *Sv2b* transcripts. **f**, Single-molecule view of a fetal human visual cortex section (Brain Image Library ace-dry-dog^74^), showing *NEFL* (green), *NELL2* (magenta) and *VIM* (yellow). Scale bar, 1 mm. **g**, *NEFL* transcripts within the region boxed in **f**, displayed as separate toggleable layers: assigned to a segmented cell (green) or unassigned (red). Left, scale bar, 100 µm; right, magnification of the boxed area, scale bar, 50 µm. **h**, Fraction of assigned and unassigned transcripts for *NEFL* (n = 4.6 million transcripts) and for all genes across the section (n = 67 million transcripts).

Within a single cell type, subcellular transcript localization can differ across genes. MERFISHEYES revealed that two glutamatergic synaptic markers, *Sv2b* (synaptic vesicle glycoprotein 2B) and *Slc17a7* (VGLUT1) in the mouse CA3 neurons, exhibit strikingly different subcellular distributions. *Sv2b* transcripts remain predominantly somatic, concentrated within the pyramidal cell layer (**Fig. 3c**). In contrast, *Slc17a7* transcripts distribute extensively into the surrounding neuropil, consistent with the established localization of mRNAs to dendritic and axonal compartments and local translation in glutamatergic circuits^58,61^. The two genes encode synaptic proteins in the same cells yet have fundamentally different mRNA localization that current cell-level analyses miss.

Finally, segmentation-based pipelines mishandle extrasomatic transcripts in two ways: some are misassigned to neighboring cells, while others remain unassigned. First, as an example, in the mouse striatum, *Gfap* is a glial gene with high expression around vascular and smooth muscle cells, whose transcripts likely localized within the large arbors of astrocytes in these regions (**Fig. 3d - blue**). Some of these transcripts are then misassigned to nearby neuronal cells (**Fig. 3d - white**) and then contaminate downstream single cell analyses. Second, in both murine and human samples there is a high level of unassigned transcripts, a loss that disproportionately affects RNAs trafficked to distal cellular processes (**Fig. 3e** - Slc17a7 higher proportion of unassigned compared to Sv2b). In fetal human visual cortex^60^ (**Fig. 3f**), we highlight NEFL, which encodes neurofilament light chain, a subunit of the neurofilament polymers that form the most abundant cytoskeletal components of axons^62^. NEFL signal extends well beyond neuronal soma into axon-rich neuropil, where densely interdigitated processes make confident assignment of transcripts to a parent cell unreliable. More broadly, 71% of the 4.6 million NEFL transcripts detected across this section fall outside segmented cell boundaries (**Fig 3g,h**). Overall, these single-molecule explorations revealed additional layers of biology regarding cell morphology and transcript localization, and highlight potential problems with current cell segmentation algorithms.

### Scalable visualization and sharing for 3D, whole-transcriptome, and million-cell datasets

We expanded MERFISHEYES beyond 2D spatial datasets to support emerging spatial transcriptomics technologies and machine learning imputations^63^ that generate 3D, whole-transcriptome, and large multi-omics data^31^. If the object uploaded contains 3D information, MERFISHEYES will automatically update the viewer to allow for 3D exploration of the volumetric organization of cell-types and gene expression patterns. As a demonstration, we visualize a 3D whole-embryo zebrafish at the 6-somite stage^31^ (**Fig. 4a**). Two cell types with well-established, contrasting anatomical positions illustrate this: the EVL/Periderm (blue), the outer epithelial monolayer that forms the embryo’s barrier with its environment^64^, and the Notochord Anterior (pink), an internal axial structure and the principal skeletal element of the early embryo^65^ (**Fig. 4b**). Viewing the same cell-types across the XY, XZ, and ZY planes reveals the precise vertical separation and volumetric alignment between these two layers that a single 2D projection cannot capture.

**Figure 4.**
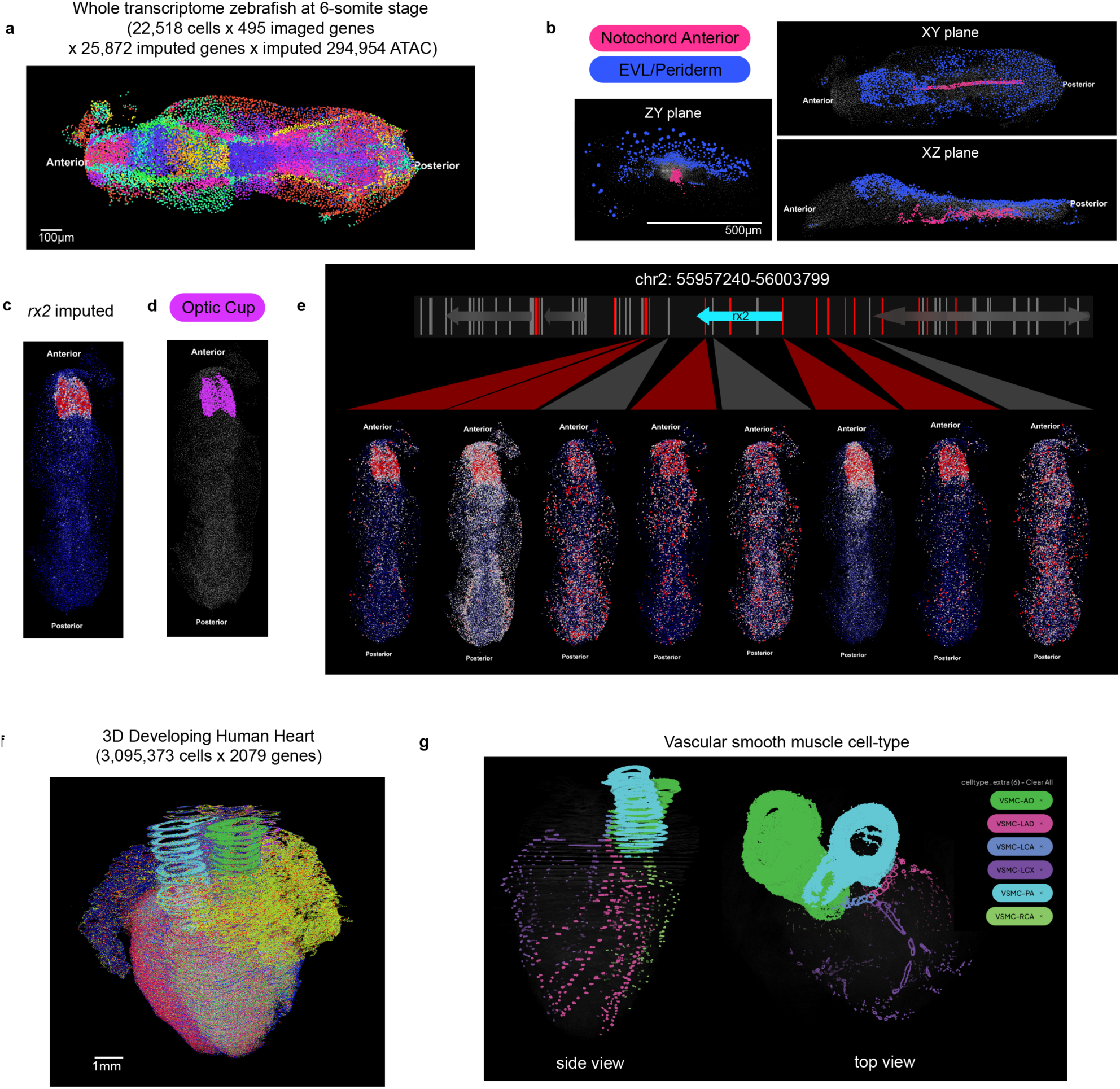
Scaling to 3D, whole-transcriptome and million-cell datasets. **a**, 3D single-cell visualization of a whole zebrafish embryo at the 6-somite stage^31^ (22,518 cells; 495 imaged genes; 25,872 imputed genes; 294,954 imputed ATAC peaks), cells colored by cell type (51 types). Anterior, left; posterior, right. Scale bar, 100 µm. **b**, Volumetric separation of two cell types in the same embryo: EVL/periderm (blue) and notochord anterior (magenta); all other cells, grey. The same selection is shown in the XY (top right), XZ (bottom right) and ZY (left) planes. Scale bar, 500 µm. **c**, Imputed *rx2* expression across the embryo (imputed counts; blue, low; red, high). **d**, The optic cup cell type (magenta) in the same embryo; all other cells, grey. **e**, Integrated genome browser at the *rx2* locus (chr2: 55,957,240-56,003,799). Top, neighbouring genes (grey arrows; *rx2*, cyan arrow) and accessible regions within 100 kb of the gene body (ticks). Bottom, accessibility of eight selected regions displayed on the embryo. Five regions (red) are accessible specifically in the optic cup, matching *rx2* expression, and are candidate regulatory elements; three (grey) show no cell-type-specific pattern. Colormap and orientation as in **c**. **f**, 3D single-cell visualization of a 12 post-conception-week developing human heart^29^ (3,095,373 cells × 2,079 genes), cells colored by cell type. Scale bar, 1 mm. **g**, Six vascular smooth muscle cell subtypes isolated from the same dataset, resolving the 3D organization of the great vessels and coronary arteries: VSMC-AO (aorta), VSMC-LAD (left anterior descending artery), VSMC-LCA (left coronary artery), VSMC-LCX (left circumflex artery), VSMC-PA (pulmonary artery) and VSMC-RCA (right coronary artery); all other cells, grey. Left, side view; right, top view. Scale bar as in **f**.

We previously integrated single-cell RNA expression and genome accessibility measurements with 3D MERFISH data within the 6-somite zebrafish embryo^31^. Specifically, the embryo shown contains 22,518 cells with 495 genes directly measured by imaging onto which the expression of 25,872 genes and 294,954 ATAC peaks were imputed from sequencing data (**Fig. 4a**). MERFISHEYES allows the imputed expression of all genes and the accessibility of their nearby regulatory elements to be displayed both spatially and in transcriptional UMAP space. We demonstrate this capability using *rx2*, a retinal homeobox gene 2, as a representative gene (**Fig. 4c**).

The imputed expression of *rx2* is restricted to the anterior “Optic Cup” population (**Fig. 4c,d**), the optic primordia at this stage, consistent with previously published in situ hybridization for rx2^66^. Selecting a gene updates a custom genome browser within MERFISHEYES showing nearby genes (arrows) and accessible genomic regions within 100 kb of the gene body. For *rx2*, we inspected eight such accessible regions; a subset (red) show accessibility confined to the same Optic Cup population, mirroring rx2 expression and marking them as candidate regulatory elements (**Fig. 4e**). Expression and accessibility can therefore be inspected side by side in the same cells of a 3D tissue, enabling identification of putative regulatory regions.

MERFISHEYES exploration scales to datasets with millions of cells, allowing for visualizing the 3D organization of entire organs. We demonstrated this with a dataset we recently produced capturing the 3D developing human heart^29^ (12 weeks post-conception) comprising over 3 million cells (3,095,373 cells × 2,079 genes) (**Fig. 4f**). By isolating the vascular smooth muscle cells (VSMC), users can resolve the 3D spatial organization of the coronary vasculature and facilitate their annotation by comparing with anatomical atlases (**Fig. 4g**). This cell-type-level navigation demonstrates that even at the scale of millions of cells, anatomically meaningful structures can be selected and examined in 3D.

Together, these examples show how MERFISHEYES scales along the two critical axes of spatial transcriptomics. Along the molecular axis, it enables whole-transcriptome and genomic-scale visualizations. Along the spatial axis, it resolves the 3D organization of millions of cells enabling exploration of organ substructures.

## Discussion

Here we introduced MERFISHEYES, a browser-based platform for exploring, comparing, and sharing single-cell and single-molecule spatial transcriptomics data. Datasets are loaded by drag-and-drop, parsed locally in the browser with no file downloads, software installation, or preprocessing, and can be shared as a link that reopens the same view on any device. At the single-cell level, the platform supports the most common analyses: the spatial organization of cell types, single-cell gene expression, cell-type abundance, and differential expression between cell types. At the single-molecule level, every detected transcript is rendered without aggregation or subsampling, preserving the subcellular signal that segmentation-based pipelines discard. This allows for checking the quality of cell segmentation and exploration of subcellular single-molecule localization in nucleus, soma, or cellular processes.

Existing spatial transcriptomics viewers, both commercial and open source, share a set of practical barriers (Table 1). They require local installation^32–34,73,74^ or self-hosted server infrastructure^38,45,69^, place preprocessing burden on the user^32,38,45,68,69^, and offer limited support for single-molecule data^33,34,37,38,69,70^. MERFISHEYES removes these barriers through three technical solutions. First, all data types are processed directly in the user’s browser into a custom compressed format with a precomputed gene index, so no preprocessing is required and a single-gene query stays instant regardless of dataset size. Files that exceed in-browser limits are converted by an equivalent server-side pipeline, so datasets of any size follow the same zero-setup workflow. Second, uploaded datasets are stored in a database-free S3 architecture that scales to millions of datasets, which removes the need for self-hosted servers and makes sharing as simple as sending a link. Third, custom rendering shaders draw more than 150 million molecules on a standard laptop, over twice the capacity of the next best browser-based tool (TissUUmaps, 70 million). This makes full-resolution, transcript-level data explorable in a browser for the first time.

MERFISHEYES serves as a visualization and data exploration infrastructure for the Brain Image Library (BIL), an NIH-funded national resource for depositing large-scale brain imaging datasets. Over the past five years, BIL has archived 158 spatial transcriptomics datasets totaling 132 million cells. However, these datasets remain difficult to access and explore: each dataset carries its own gene library, cell-type definitions, and segmentation pipeline, so comparisons between datasets are unsubstantiated. We mapped all deposited datasets to a common reference taxonomy and made them available in MERFISHEYES, turning an archive into an explorable resource. This provides the first interactive single-cell and single-molecule access to over a hundred brain datasets spanning multiple species.

Hypotheses can now be tested directly against these published data, as we demonstrated with three examples: (1) identifying spatially differential expressed genes within specific neuronal subclasses populations, that are reproducible across spatial datasets, but are not captured within scRNA-seq data, (2) cell-type specific quantification of gene expression changes in neurodegenerative mouse models (ex: *Apoe* upregulation in microglia in Alzheimer’s disease model), and (3) exploring subcellular localization of individual transcripts revealing that different genes are preferentially localized.

Collapsing transcript positions into cell labels loses several kinds of information. The single-molecule view allows for visualizing and comparing cell morphology across cell-types, i.e. striatal *ChAT*+ interneurons are ∼2 fold larger compared to medium spiny neurons across all species. Second, transcripts across genes can have different subcellular organization, i.e. *Sv2b* has more restricted somatic expression compared to *Slc17a7*, which is potentially trafficked to processes of hippocampal pyramidal neurons. Third, the single-molecule view allows for inspecting misassignment of transcripts and discarded transcripts outside of cell segmentation, i.e. *Gfap* transcripts expressed in non-neuronal cells are sometimes misassigned to neighboring neurons and transcripts outside segmentation masks are entirely omitted in downstream cell-level analysis. For example, 71% of NEFL transcripts in the fetal human visual cortex are dismissed in the downstream cell-level analyses. Recent frameworks have begun analyzing rather than discarding these transcripts by classifying RNA in cellular protrusions and clustering extrasomatic transcripts into synaptic, dendritic, and axonal RNA granules^71,72^. MERFISHEYES complements them on the visualization side: molecules outside the segmentation are displayed as separate toggleable layers, so the extent and organization of this signal can be inspected rather than dismissed.

MERFISHEYES has two main limitations, which define our future development. First, the platform does not yet display image-based layers such as nuclear (DAPI), protein, or chromatin channels, or cell-segmentation boundaries (Table 1). We plan to add these layers so that transcripts can be inspected directly against their tissue and cellular context. Second, visualizing imputed whole-transcriptome and multiome data requires custom processing; we aim to build imputation directly into the platform, open the multiome pipeline to user uploads, and extend integration to additional single-cell and single-nucleus modalities such as DNA accessibility (ATAC) and DNA methylation.

As spatial transcriptomics grows in scale and modality, we envision MERFISHEYES will continue to support data exploration and visualization as an easy to access web-based open-source resource available at merfisheyes.com (github.com/kresnajenie/merfisheyes).

## Methods

### Datasets

No new data were generated for this study. Every dataset shown is publicly available through the Brain Image Library (BIL), the Allen Institute or the original publication, and was used as deposited unless stated otherwise below. Supplementary Table 1 lists each dataset shown in a figure with its source publication, platform, species, cell and transcript counts, BIL accession and the processing applied here.

### Web application architecture

MERFISHEYES is a client-side single-page application built with Next.js 16.0.7 and React 18.3.1 in TypeScript 5.6, styled with Tailwind CSS 4.1.11 and HeroUI 2.8.4, and served as a static bundle from Cloudflare. Client state is held in Zustand 5.0.8. Parsing, indexing and rendering all run in the user’s browser. Cell-level and molecule-level data leave the device only when the user explicitly uploads a dataset, as described below.

Two data paths are kept separate. In the local path, files selected by drag-and-drop are read through the browser File API, parsed inside a Web Worker and held in page memory; closing the tab discards them. In the shared path, started only by a user action, the parsed dataset is re-encoded into the chunked format below and transferred to cloud object storage. The raw source files are never uploaded on this path.

A small server component handles sharing and the dataset catalogue: Next.js API routes backed by PostgreSQL through Prisma 6.17, authentication with NextAuth v5 (Google OAuth), object storage through the AWS SDK for JavaScript v3 (@aws-sdk/client-s3, @aws-sdk/s3-presigned-post, @aws-sdk/s3-request-presigner) and notification email through Amazon SES.

Rendering requires WebGL2. Dataset ceilings in the local path are set by the browser’s per-tab memory limit, not by MERFISHEYES: roughly 2-4 GB on Chrome and Firefox and 2-3 GB on Safari, with 64-bit maxima near 8 GB (Chrome, Firefox) and 4 GB (Safari), and a per-file limit of about 2 GB.

Viewer state is serialized into the URL so that a link reproduces a view. State is encoded as compact JSON, base64url-encoded and carried in the query string; the left and right datasets of a split view occupy the v and rv keys. For single-cell views the state holds the dataset, selected gene and metadata column, selected cell types, visualization mode, gene and numerical scale bounds, point-size scale, embedding, column-type overrides, 2D/3D mode, colormap, scene rotation and axis flips, and, for two-gene co-expression, the second gene with its scale and swap state. For single-molecule views it holds the per-gene tuples (gene, colour, local scale, visibility, assigned/unassigned display flags), the global scale and the view mode. Camera position and target are not serialized, so a shared link restores the visualization state but not the viewpoint.

### Supported input formats and in-browser parsing

Single-cell data are accepted as MERSCOPE output folders, Xenium output folders, AnnData .h5ad files and MERFISHEYES chunked folders. Single-molecule data are accepted as .csv, .parquet and MERFISHEYES chunked folders.

Format-specific adapters (H5adAdapter, H5adZarrAdapter, XeniumAdapter, MerscopeAdapter, ProcessedSingleMoleculeAdapter, ChunkedDataAdapter) each emit a common in-memory representation (StandardizedDataset for single-cell data, SingleMoleculeDataset for single-molecule data). HDF5-backed .h5ad files are read with h5wasm 0.8.6 and anndata.js 0.0.2; Zarr-backed AnnData with zarrita 0.5.1; Parquet with hyparquet 1.20 and hyparquet-compressors 1.1.1; delimited text with PapaParse 5.5.3 in chunked streaming mode; gzip streams with pako 2.1.

Parsing runs in dedicated Web Workers (standardized-dataset.worker, single-molecule.worker, zarr-densify.worker) that talk to the main thread through Comlink 4.4.2, so file I/O never blocks the render loop. Coordinates are stored as Float32Array; expression is stored sparsely. Negative-control probes, unassigned codewords and blank barcodes are removed from single-molecule datasets before display.

### Compressed chunked format

Uploaded datasets are re-encoded into a chunked, gzip-compressed layout designed for range-request streaming:

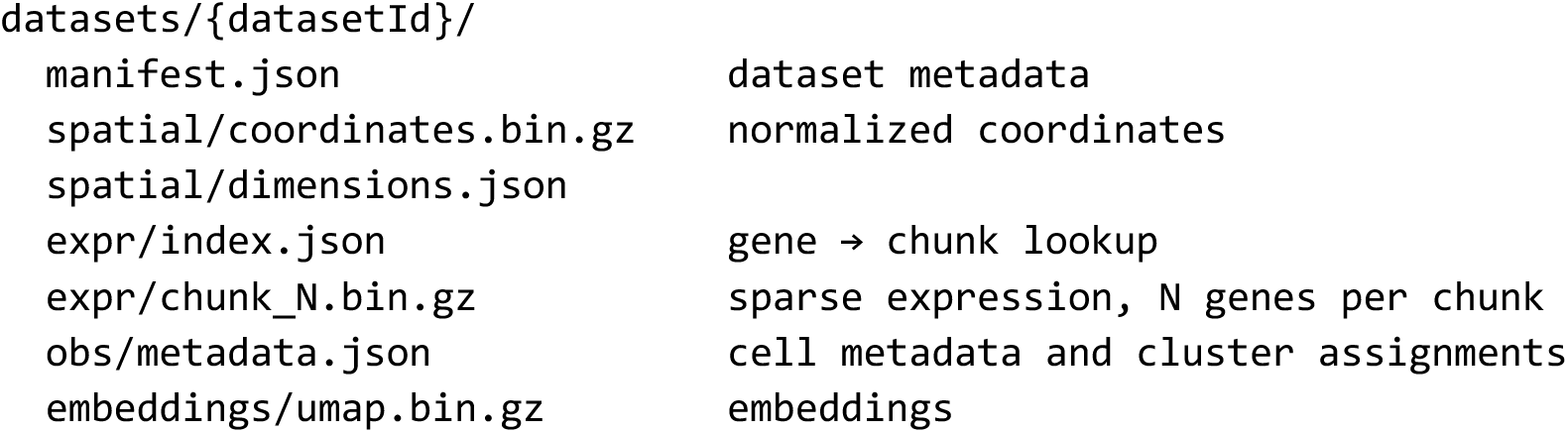

Each expression chunk starts with a 16-byte header (version, num_genes, chunk_id, total_cells; all uint32), then a 24-byte-per-gene table (gene_offset uint64, gene_length uint64, nnz uint32, reserved uint32), then per-gene sparse payloads as a uint32 array of cell indices and a parallel float32 array of values. Sparse encoding cuts expression storage by about 90% on typical panels.

Genes per chunk depend on panel size: 50 below 100 genes, 100 below 500, 200 below 2,000, 500 below 10,000 and 1,000 above. At load time the viewer fetches only manifest.json and expr/index.json. Expression chunks are fetched when a gene is selected, decompressed with the browser’s native DecompressionStream and cached in the worker for other genes in the same chunk.

### Client-side conversion, upload and link sharing

On Upload & Save, the parsed dataset is encoded into the chunked format in the browser. A content fingerprint is computed first as a SHA-256 digest over dataset structure (cell count, gene count, dataset type) plus sampled content (every third gene name and every fifth cell coordinate), excluding file names and timestamps, so the same data uploaded twice is recognized whatever it was called. The fingerprint is checked against the catalogue (/api/datasets/check-duplicate/{fingerprint}); if the dataset exists, its link is returned instead of re-uploading.

Otherwise the client requests an upload session (/api/datasets/initiate), receives presigned POST credentials scoped to that dataset prefix, uploads each chunk directly to object storage without passing through the application server, and marks the transfer complete (/api/datasets/{datasetId}/files/{fileKey}/complete, then /api/datasets/{datasetId}/complete). Single-molecule datasets use a parallel set of routes under /api/single-molecule/.

Converted datasets are stored in Amazon S3. The viewer reads them through presigned URLs issued per dataset by /api/datasets/{datasetId} and fetched lazily by the worker. Each dataset has a persistent identifier that resolves to merfisheyes.com/viewer/{id}. On completion the user receives the link and dataset metadata (cell count, gene count, platform) by email through Amazon SES.

### Server-side conversion for large datasets

Datasets that exceed in-browser limits can be processed on a server instead, so the browser only transfers bytes and does no parsing. This path is available to signed-in users.

Raw files are uploaded unmodified to raw/{datasetId}/ in S3 with relative paths preserved, so Xenium and MERSCOPE folders arrive intact. Files below 100 MB use a single presigned PUT; larger files use S3 multipart upload with 32 MB parts, presigned per part and uploaded in parallel by the client, which lifts the ∼5 GB single-PUT ceiling. Per-file and session completion are recorded through /api/ingest/{id}/files/{key}/complete and /api/ingest/{id}/complete; an in-flight upload can be cancelled through /api/ingest/{id}/abort, which aborts the multipart upload and deletes any objects already written.

On session completion the application submits a job to AWS Batch, which runs the processing container on ECS Fargate (us-west-2). Job resources come from the user’s compute tier: standard, 4 vCPU / 16 GiB; large, 8 vCPU / 32 GiB; xlarge, 16 vCPU / 64 GiB. Jobs that run out of memory are retried automatically on the next tier.

The container is built on python:3.12-slim with pinned dependencies (numpy 2.2.6, pandas 2.3.3, scipy 1.14.1, anndata 0.12.10, h5py 3.16.0, pyarrow 18.1.0, boto3 1.42.85) and runs the same process_spatial_data.py and process_single_molecule.py used for local conversion, so the browser, command-line and server paths all write the same chunked layout from one implementation. The entrypoint downloads the raw prefix to Fargate ephemeral storage, runs the processor, uploads the result to datasets/{datasetId}/ and reports status. Raw uploads are deleted from object storage on success, so no unprocessed user data is kept. Cell-type annotation is not run on the server; users can supply a per-cell label CSV, which is merged during chunking.

Status is reported to the application through an HMAC-signed callback (/api/ingest/{id}/callback) rather than by writing to the database from the worker, so database credentials stay in the application. Datasets move through UPLOADING → QUEUED → PROCESSING → COMPLETE or FAILED; the worker maps the processors’ staged log output (reading, coordinates, expression chunks, DE statistics, upload) onto progress updates, posted best-effort and abandoned after three consecutive failures. /api/ingest/{id}/status is a polling fallback. On completion the user is emailed a viewer link.

Compute cost per dataset ($0.01-0.30) was estimated from actual AWS charges at September 2026 Fargate and S3 pricing, for datasets with a final compressed size of about 300 MB.

### Command-line conversion utility

The chunked format can also be produced outside the browser with scripts/process_spatial_data.py (single-cell) and scripts/process_single_molecule.py (single-molecule).

process_spatial_data.py takes an .h5ad file or a Xenium/MERSCOPE folder and an output directory, with --format {h5ad,xenium,merscope} to override format detection, --chunk-size to override the genes-per-chunk heuristic, --workers for parallel chunk writing, --mask / --mask-col / --mask-keep to drop cells by a boolean metadata column (default is_artifact), --mmc-csv to merge MapMyCells assignments, --reorder, and --de-max-celltypes (default 20,000) to bound the differential-expression precomputation.

process_single_molecule.py takes a Parquet or CSV file, or a directory of per-sample subfolders, and an output folder, with --dataset-type {xenium,merscope,custom} selecting column mappings and --gene-col / --x-col / --y-col / --z-col / --cell-id-col overriding them. Molecules whose cell-assignment value is −1 are treated as unassigned and written to separate files. --s3-prefix and --mapping-url write the mapping file that links a single-cell dataset to its single-molecule counterpart, --link-column (default _sample_id) names the cluster column the viewer reads for that link, and --workers / --sample-workers control parallelism within and across samples.

### Rendering

Points are drawn with Three.js 0.180 as one THREE.Points object per layer, backed by a BufferGeometry whose position, color, size and alpha attributes are Float32Array buffers written in place; parsed coordinate buffers are handed to the geometry without an intermediate copy. All points in a layer share one draw call and one geometry, so cost scales with buffer size rather than object count, which is what lets the full molecule set be drawn without subsampling.

A custom GLSL shader pair replaces the default point material. The vertex shader sets point size in world space as gl_PointSize = size * dotSize * projectionMatrix[1][1] / - mvPosition.z, clamped to [0.5, 200] pixels, where projectionMatrix[1][1] equals 1/tan(fov/2); points therefore scale with perspective, and a per-point size attribute carries expression- or slider-driven modulation. The fragment shader discards fragments beyond a radius of 0.5 in point-coordinate space and applies a smoothstep falloff between 0.5 and 0.45, giving anti-aliased circular points.

The renderer is a THREE.WebGLRenderer with antialias: true and preserveDrawingBuffer: true (so the canvas can be read back for figure export), sized to the container at window.devicePixelRatio. The camera is a THREE.PerspectiveCamera with a 75° vertical field of view, near and far planes at 0.1 and 10,000, at z = 500 on a black background.

Camera interaction depends on dimensionality. In 2D mode the viewer uses OrbitControls with rotation disabled and mouse bindings remapped so that left-drag pans, middle-drag dollies and right-drag rotates, with damping at 0.25. In 3D mode, which is enabled automatically when a dataset carries a third coordinate, the viewer uses TrackballControls with unconstrained rotation and rotation, zoom and pan speeds of 1.5, 1.2 and 0.8, with dynamic damping of 0.15. A requestAnimationFrame loop updates the controls and re-renders each frame; a ResizeObserver tracks container size so split panes rescale correctly.

### Rendering-capacity benchmark

The maximum point count in Fig. 1f was measured on a MacBook Air M3 in Google Chrome. For MERFISHEYES, single-molecule data were streamed into a single view until the tab failed; the reported value is the largest count that rendered and remained interactive. For each comparison tool, the same data were loaded at increasing point counts until the tool crashed, froze or refused the file. Tools, versions, access dates and per-tool outcomes are given in Table 1.

### Brain Image Library integration

BIL datasets are catalogued in a PostgreSQL database and served to merfisheyes.com/explore through /api/explore, which returns only published, non-internal entries with a resolvable storage location. The endpoint supports free-text search and filtering by species, tissue, platform, gene and dataset type, with pagination. Conversion of BIL datasets to the chunked format runs as a SLURM pipeline on the PSC Bridges-2 cluster (scripts/bil-scripts/), with separate job scripts for slice combination, MapMyCells mapping, single-cell and single-molecule processing, and synchronization to object storage.

### Cell-type standardization with MapMyCells

Cell-type labels were assigned with the Allen Institute cell_type_mapper package through its FromSpecifiedMarkersRunner (cell_type_mapper.cli.from_specified_markers), called by scripts/map_my_cell.py. Mouse datasets were mapped against the ABC whole-mouse-brain reference (precomputed statistics precomputed_stats_ABC_revision_230821.h5, marker set mouse_markers_230821.json, drop level CCN20230722_SUPT). Human and rhesus macaque datasets were both mapped against the Siletti human reference (precomputed_stats.siletti.training.h5, markers query_markers.n10.20240221800.json, drop level CCN202210140_SUPC); marmoset datasets were not mapped. Input normalization was set to raw. Gene identifiers were harmonized to the reference through the per-species gene.csv mapping. Mapping ran on the PSC Bridges-2 cluster, with n_processors and the memory ceiling set from the available cores and RAM.

### Assessment of mapping quality

For each dataset, mapping quality (Supplementary Fig. 2a) is the mean over all cells of the MapMyCells average bootstrap correlation at the cluster level, as reported in the cell_type_mapper output. Group means are reported in the figure legend. Mouse datasets from neonatal animals are shown separately because they map poorly to the adult reference. The same per-dataset value is shown on the Explore page as a guide to annotation quality.

### Split-synchronized comparison view

Two datasets can be shown side by side in a resizable split pane, with the left and right datasets carried in the URL as the v and rv parameters, each with its own visualization state, so a comparison is itself shareable as a link. Selection state is synchronized in both directions: the active metadata column, selected and hidden cell types, selected gene, colour palette, and the co-expression gene and swap state. Changing a gene or cell-type selection on one side applies the same choice to the other. Camera position is not synchronized; each pane is navigated independently.

### Differential expression and co-expression

During parsing, MERFISHEYES precomputes per-cell-type summaries for every gene: mean expression, the fraction of cells with non-zero expression, and per-cell-type cell counts, from which global per-gene sums and fold change versus a reference population are derived without a second pass over the matrix. Summaries are computed on the matrix as stored by the adapter, raw or normalized, with no further transformation. The cell-type column is chosen automatically as the first categorical column found in the order class_name, subclass_name, supertype_name, cluster_name, subcluster_name, super_cluster_name, leiden, Cluster.

In the two-gene co-expression mode, each cell is coloured by two independent colour ramps (Fig. 2d,e: *Pdyn*, green; *Drd1*, magenta), each scaled to the bounds set in the interface; cells high in both appear white by additive mixing.

### Genotype comparison of microglial Apoe expression

Microglia were selected as the MapMyCells subclass in each section of the four-genotype cohort^55^. Raw *Apoe* counts per microglial cell were pooled across sections within a genotype (4, 4, 5 and 5 sections for WT, Trem2R47H, Trem2R47H;5xFAD and 5xFAD) and plotted by genotype (Fig. 2i). The comparison is descriptive; no hypothesis test was applied.

### Soma size from single-molecule clustering

Somata were reconstructed from transcript positions alone. *ChAT* transcripts (cholinergic interneurons) and *Drd1* transcripts (D1 medium spiny neurons) in striatal sections from rhesus macaque^60^ and mouse^50^ were clustered as follows. Single-molecule spots (assigned and unassigned combined) were taken in raw micron coordinates and reduced to 2D (x, y; sections are ∼6 µm thick), and restricted to a striatum region drawn by hand in napari (point-in-polygon test). Spots were clustered with DBSCAN (scikit-learn; Euclidean distance in µm) with min_samples = 15 and eps = 10 µm for *ChAT* in both species. For *Drd1*, eps was set per dataset (7 µm macaque, 6 µm mouse) as the value that resolved the largest number of clusters before neighbouring D1 somata merged or fragmented; a single eps suffices for the sparse cholinergic cells but not for densely packed D1 cells. Noise points (label −1) were discarded.

For each cluster with at least three points, the effective soma diameter was computed as 2√(A/π), where A is the area of the 2D convex hull (scipy.spatial.ConvexHull). In mouse, clusters were retained only if their centroid lay within 25 µm (nearest neighbour) of a single-cell centroid annotated by MapMyCells as 058 PAL-STR Gaba-Chol (for *ChAT*) or 061 STR D1 Gaba (for *Drd1*). In macaque, where matching single-cell labels were not available, *ChAT* clusters with at least 60 transcripts and all *Drd1* clusters were retained. Diameters are summarised as median and interquartile range, compared within species by a one-sided Mann–Whitney U test (*ChAT*+ > D1), and reported as the ratio of medians. Because MERFISH+ and MERSCOPE differ in detection sensitivity, absolute diameters are not comparable across platforms; only the within-species ratio is interpreted.

### Transcript assignment analysis

For each single-molecule dataset, a transcript was counted as assigned if it carried a cell identifier and unassigned otherwise (−1 or empty in the vendor cell column). Fractions (Fig. 3e,h) were computed over the whole section after removing blank barcodes and negative controls. For Fig. 3d, cell centroids were labelled by MapMyCells class: striatal neurons in white, and astrocytes, oligodendrocytes, microglia, vascular and muscle cells in blue.

### 3D visualization, imputed whole-transcriptome and chromatin accessibility

When an uploaded dataset carries a third spatial coordinate, the viewer switches to 3D mode with the trackball camera described above and adds XY, XZ and ZY projections.

Imputed gene expression and ATAC peak accessibility for the 6-somite zebrafish embryo (22,518 cells) were taken directly from Wan et al. 2026^31^, where 25,872 genes and 294,954 peaks were imputed from single-cell RNA-seq and ATAC-seq onto the 495-gene MERFISH measurement with Tangram^63^. No imputation was re-run here. Imputed values are displayed as counts on the same colour ramp as measured genes.

The integrated genome browser reads a BED file of peak coordinates and gene annotations (GRCz11). Selecting a gene displays neighbouring genes and all accessible regions within 100 kb on either side of the gene body; selecting a region colours cells by its imputed accessibility. Regions called spatially significant in Fig. 4e are those reported as cell-type-associated in Wan et al. 2026; no new scoring was done here.

### Processing-time benchmark

Processing times in Supplementary Fig. 1 were measured on a databank of 56 datasets (23 Xenium, 25 MERSCOPE and 8 MERFISH runs; 48 with single-cell and 34 with single-molecule tables) drawn from 10x Genomics Xenium demonstration data (7), Vizgen MERSCOPE showcase data (20), Brain Image Library deposits (21) and unpublished mouse brain MERFISH data from this laboratory (8); identifiers, sizes and sources are listed in Supplementary Table 2. Inputs (Xenium, MERSCOPE, .h5ad and single-molecule tables) were each processed once through the browser path and once through the server path. Browser runs used Google Chrome [version] on a workstation with 64 cores and 503 GB RAM; per-file and per-tab limits are set by the browser and match those on a laptop. Server runs used the default 4 vCPU / 16 GiB tier, with automatic retry on 32 or 64 GiB where noted in the figure. Times cover parsing and conversion only and exclude network transfer and job queueing. Datasets that the browser refused (over ∼2 GB), that crashed or froze the tab, or whose whole-transcriptome matrix was skipped are marked in the figure. Cell count, gene count, metadata columns and chunk count were checked to match between the browser and server outputs for every dataset that both completed.

### Statistics and reproducibility

Soma diameters (Fig. 3b) were compared within species by a one-sided Mann-Whitney U test; n somata per group are given in the legend. All other comparisons in the manuscript are descriptive and no other statistical tests were applied. Each benchmark measurement (Supplementary Fig. 1) is a single run.

## Data availability

All datasets analysed are publicly available and are cited in the References; Brain Image Library accessions and Allen Institute sources for each figure dataset are listed in Supplementary Table 1. Every dataset shown is also viewable in MERFISHEYES at the links given in Supplementary Table 1.

## Code availability

MERFISHEYES source code, the conversion scripts and the BIL processing pipeline are available at https://github.com/kresnajenie/merfisheyes.

## Notes

https://www.merfisheyes.com

### Competing Interest Statement

The authors have declared no competing interest.

### Summary of Updates

I am revising the headers of my manuscript pdf. It previously contained private information.

