## Supplementary table and figures for "MERFISHEYES: Web-based Visualization and Sharing Platform for Single-cell and Single-molecule Spatial Transcriptomics"

Supplementary Table 1 | Datasets shown in this study.

| Dataset | Shown in | Platform | Species | Cells / transcripts | Source | Accession |
|---|---|---|---|---|---|---|
| Allen Whole Mouse Brain Atlas | Fig. 1a-c; Fig. 3c,d | MERSCOPE | mouse | ~4 M cells | [47] | ABC Atlas MERFISH-C57BL6J-638850; BIL ace-den-fix (https://api.brainimagelibrary.org/web/view?bildid=ace-den-fix) |
| P28 developing mouse brain, posterior | Fig. 2a-c,e; Sup. Fig. 2c; Fig. 3a,b (mouse) | MERSCOPE | mouse | 125k cells | [50] | BIL BIL ace-low-and (https://api.brainimagelibrary.org/web/view?bildid=ace-low-and) (https://api.brainimagelibrary.org/web/view?bildid=ace-low-and) |
| P28 developing mouse brain, anterior | Fig. 2d | MERSCOPE | mouse | 140k cells | [53] | BIL ace-low-bag (https://api.brainimagelibrary.org/web/view?bildid=ace-low-bag) |
| Four-genotype AD cohort (WT, Trem2R47H, Trem2R47H;5xFAD, 5xFAD; 12 months) | Fig. 2f-i | MERSCOPE | mouse | 18 sections; 3 million cells | [55,73] | BIL ace-ear-nap (https://api.brainimagelibrary.org/web/view?bildid=ace-ear-nap) |
| Adult rhesus macaque subcortical atlas | Fig. 3a,b (macaque) | MERSCOPE | rhesus macaque | 3 million cells | [60] | BIL ace-dud-wag (https://api.brainimagelibrary.org/web/view?bildid=ace-dud-wag) |
| Mouse striatum, GFAP misassignment | Fig. 3d | MERSCOPE | mouse | 125k cells | [50] | ace-low-and |
| Fetal human visual cortex (GW24) | Fig. 3f-h | MERSCOPE | human | 67 million transcripts | [74] | BIL ace-dry-dog (https://api.brainimagelibrary.org/web/view?bildid=ace-dry-dog) |
| 6-somite zebrafish embryo, multiome | Fig. 4a-e | MERFISH + imputation | zebrafish | 22,518 cells; 495 imaged genes; 25,872 imputed genes; 294,954 ATAC peaks | [31] | [deposit accession for Wan et al. 2026] |
| 12 pcw developing human heart | Fig. 4f,g | MERFISH+ | human | 3,095,373 cells × 2,079 genes | [29] | [deposit accession for Kern et al. 2025] |
| Benchmark datasets (Sup. Fig. 1) | Sup. Fig. 1 | Xenium, MERSCOPE, .h5ad, single-molecule | Mouse, macaque, human | listed in figure | 56 datasets: 10x Genomics Xenium demonstration datasets (7), Vizgen | Public download pages of 10x Genomics and Vizgen; BIL IDs as listed under Source |

| | | | | | |
|---|---|---|---|---|---|
| | | | | | MERSCOPE showcase datasets (20), Brain Image Library deposits (21: 16 mouse spinal cord Xenium sections ace-pie-*, ace-irk-sag, ace-dry-dog, ace-ear-nap, ace-dip-tub) and unpublished mouse brain MERFISH datasets from this laboratory (8); sample identifiers in Supplementary Table 2 |

All repository links were accessed on 9 September 2026. Brain Image Library deposits are unversioned and were used as deposited; the Allen ABC Atlas release used is given in the Accession column.

### Supplementary Figures

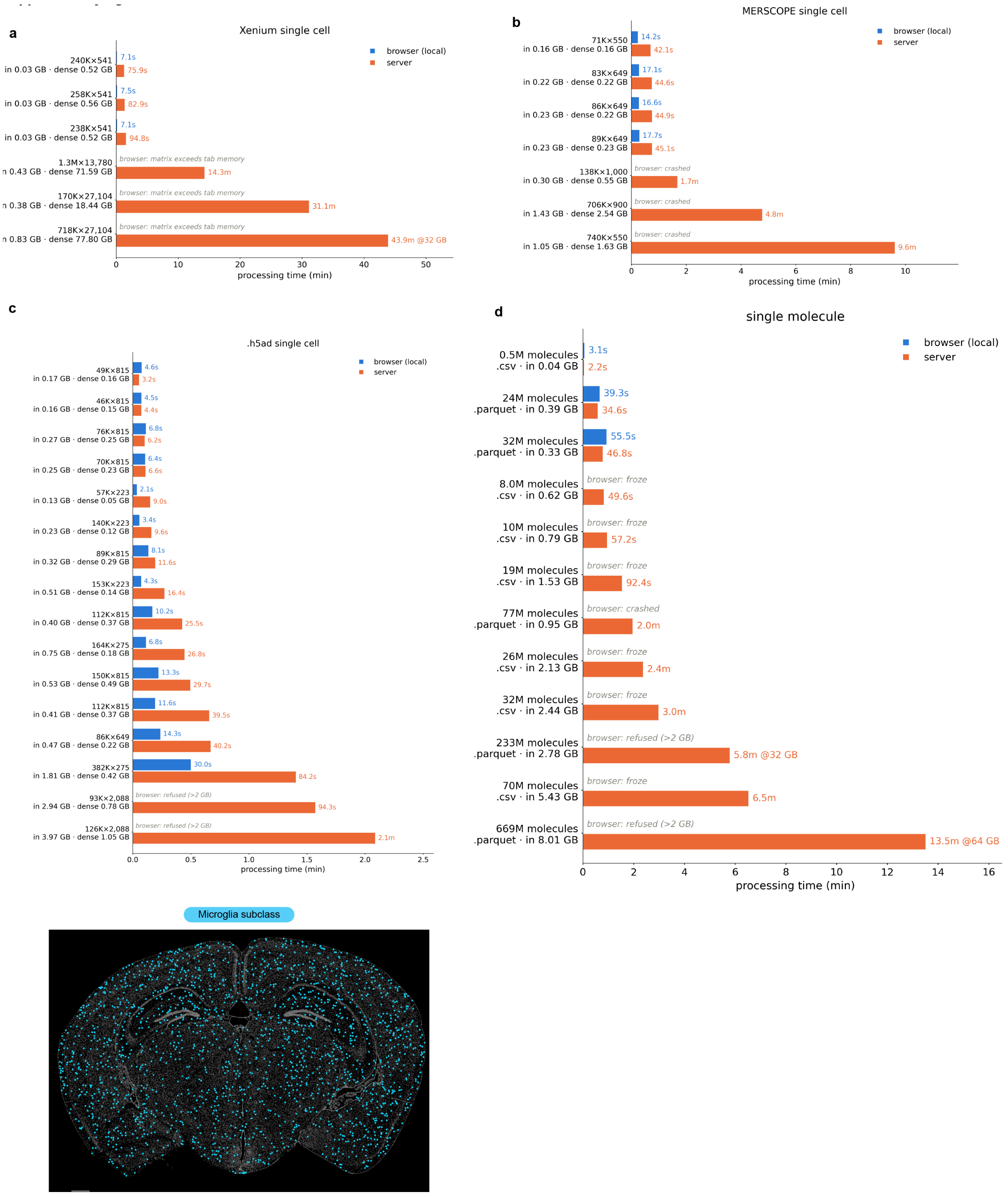

**Supplementary Figure 1 | Browser and server processing times for real datasets, and a broadly distributed cell type. a-d**, Conversion time for the same dataset processed in the browser (blue) and server-side (orange), for (**a**) Xenium, (**b**) MERSCOPE and (**c**) .h5ad single-cell inputs, and (**d**) single-molecule inputs. Each dataset is labelled with its dimensions (cells × genes, or transcript count and file format), the input size read by the pipeline and, for single-cell data, the size of its expression matrix in memory (dense, float32). Datasets are ordered by processing time, which tracks dense matrix size. Grey italics mark datasets the browser could not process: files over the ~2 GB per-file limit were refused, tabs that exceeded their memory cap crashed or froze, and whole-transcriptome panels (18-78 GB dense) loaded cells and gene lists but skipped the expression matrix. Server jobs that exceeded the default 16 GB instance were retried automatically on a larger one; their labels give the instance that completed them (for example, 43.9 m @32 GB). Each bar is a single run. Datasets were drawn from public Xenium demonstration datasets (10x Genomics), public MERSCOPE datasets (Vizgen), Brain Image Library deposits (including 16 mouse spinal cord Xenium sections, ace-pie-*) and unpublished mouse brain MERFISH datasets from this laboratory; identifiers and sources are listed in Supplementary Table 2. Browser measurements were made in Chrome 137.0.7151.103 on a workstation with 64 cores and 503 GB RAM; the limits are set by the browser tab, not the hardware, and the per-file and heap caps are the same as on a laptop. Times exclude network transfer and job queueing. **e**, Single-cell view of the microglia subclass (cyan) across the coronal section of the Allen Institute Whole Mouse Brain Atlas shown in Fig. 1a; all other cells, grey. Unlike the layer-restricted neocortical subclasses in Fig. 1a, microglia are distributed throughout the section. Scale bar, 500 µm.

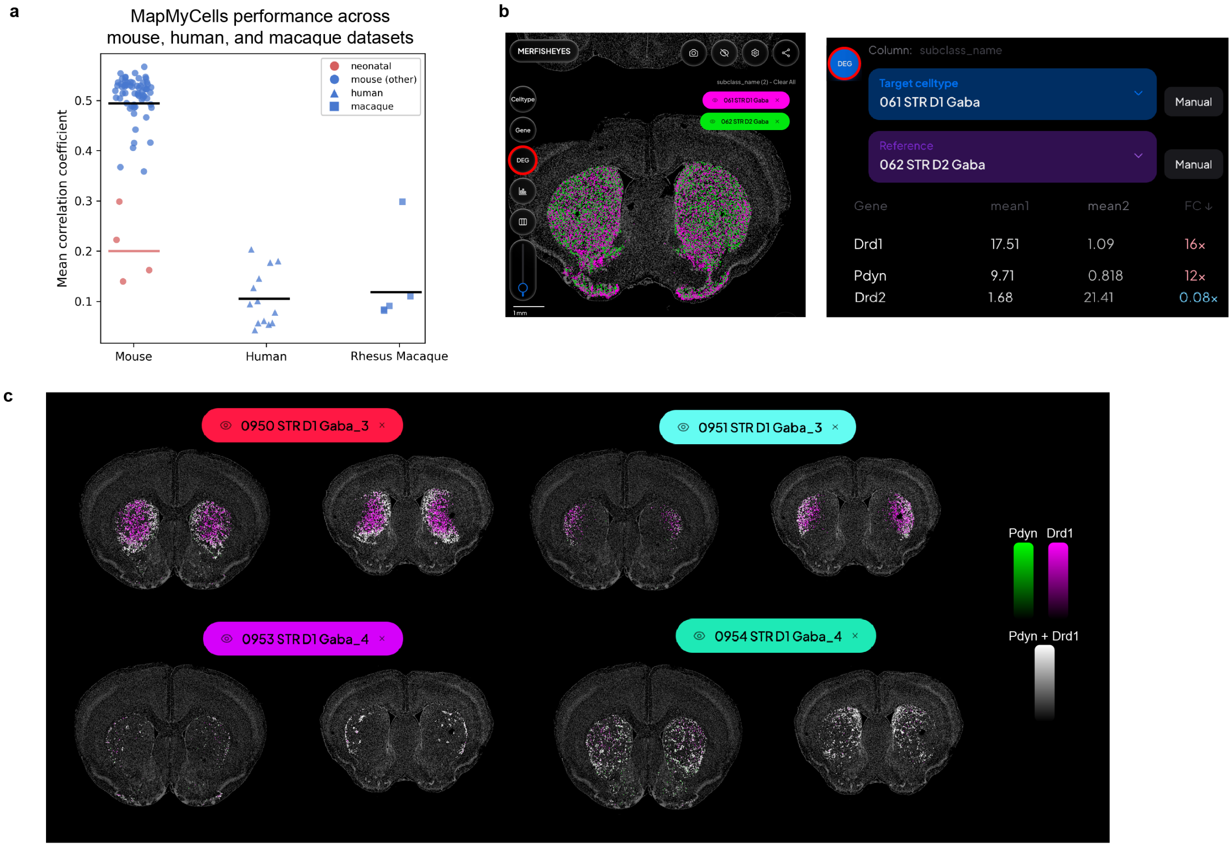

**Supplementary Figure 2 | Cell-type mapping quality and cluster-level reproducibility. a**, MapMyCells mapping quality for every mouse, human and rhesus macaque dataset in the Brain Image Library. Each point is one dataset; the value is MapMyCells' average correlation to the Allen Institute reference taxonomy over bootstrap runs. Horizontal lines are group means: mouse, 0.491 (n = 68 datasets, of which the 4 neonatal datasets are shown in red, mean 0.206; the 64 non-neonatal datasets have mean 0.509); human, 0.106 (n = 13); rhesus macaque, 0.133 (n = 5). **b**, The MERFISHEYES differential expression interface, comparing the 061 STR D1 Gaba subclass (target) against 062 STR D2 Gaba (reference) in the ABC dataset. Genes are ranked by fold change; the table reports mean expression in the target and reference populations and their ratio. **c**, *Pdyn* (green) and *Drd1* (magenta) co-expression shown separately for four D1 clusters: 0950 STR D1 Gaba_3, 0951 STR D1 Gaba_3, 0953 STR D1 Gaba_4 and 0954 STR D1 Gaba_4. Within each pair, left is the P28 dataset and right is the ABC atlas, at the same anterior striatal section for both. Cells expressing both genes appear white.
